# Non-invasive laminar inference with MEG: Comparison of methods and source inversion algorithms

**DOI:** 10.1101/147215

**Authors:** James J Bonaiuto, Holly E Rossiter, Sofie S Meyer, Natalie Adams, Simon Little, Martina F Callaghan, Fred Dick, Sven Bestmann, Gareth R Barnes

## Abstract

Magnetoencephalography (MEG) is a direct measure of neuronal current flow; its anatomical resolution is therefore not constrained by physiology but rather by data quality and the models used to explain these data. Recent simulation work has shown that it is possible to distinguish between signals arising in the deep and superficial cortical laminae given accurate knowledge of these surfaces with respect to the MEG sensors. This previous work has focused around a single inversion scheme (multiple sparse priors) and a single global parametric fit metric (free energy). In this paper we use several different source inversion algorithms and both local and global, as well as parametric and non-parametric fit metrics in order to demonstrate the robustness of the discrimination between layers. We find that only algorithms with some sparsity constraint can successfully be used to make laminar discrimination. Importantly, local t-statistics, global cross-validation and free energy all provide robust and mutually corroborating metrics of fit. We show that discrimination accuracy is affected by patch size estimates, cortical surface features, and lead field strength, which suggests several possible future improvements to this technique. This study demonstrates the possibility of determining the laminar origin of MEG sensor activity, and thus directly testing theories of human cognition that involve laminar- and frequency- specific mechanisms. This possibility can now be achieved using recent developments in high precision MEG, most notably the use of subject-specific head-casts, which allow for significant increases in data quality and therefore anatomically precise MEG recordings.

## 1 Introduction

Modern theories of brain organization and function increasingly incorporate the laminar organization of cortical projections and oscillatory signatures of neural activity (Adams et al., 2013; Arnal and Giraud, 2012; Bastos et al., 2012; Jensen et al., 2015; Wang, 2010). While high resolution functional magnetic resonance imaging (fMRI) can resolve laminar-specific activity (Chen et al., 2013; Goense et al., 2012; Guidi et al., 2016; Huber et al., 2015; Kok et al., 2016; Koopmans et al., 2011, 2010; Olman et al., 2012; Scheeringa et al., 2016), it cannot measure dynamics at a millisecond time-scale. Being a completely non-invasive and direct measure of neuronal activity capable of such temporal resolution, magnetoencephalography (MEG), is an attractive option for testing such theories (Baillet, 2017). While MEG has excellent temporal resolution, its spatial resolution is limited by subject movement and co-registration error (Hillebrand and Barnes, 2011, 2003; Medvedovsky et al., 2007; Uutela et al., 2001). With recently developed high precision MEG using head-cast technology, it is possible to address both issues and record higher quality MEG data than previously achievable (Liuzzi et al., 2016; Meyer et al., 2017; Troebinger et al., 2014a, 2014b). Simulations have shown that such high quality data make it theoretically possible to distinguish the MEG signal originating from either deep or superficial laminae (Troebinger et al., 2014a). However, previous attempts at laminar localization have used global measures of model fit that cannot infer the laminar origin of activity in a spatially specific way, using regions of interest (ROIs). Additionally, multiple source inversion algorithms exist to link extra-cranial electromagnetic activity measured at the sensors to cortical sources. Each of these algorithms uses different assumptions to constrain the source estimation, further complicating the validation of laminar specific MEG by making it unclear which might be most suited for laminar-specific analysis.

We here develop whole brain and ROI analyses using multiple source inversion algorithms for classifying MEG signals as originating from either deep or superficial laminae, test them using simulated laminar data, and compare them in terms of their classification performance. To this end, we simulated sensor data for source activity at locations on either the pial or white matter cortical surface (representing superficial and deep cortical laminae, respectively). We then measured the accuracy of two types of analyses (whole-brain or ROI) in determining, based on the sensor data alone, the correct origin of the source. The whole brain analysis reconstructs the sensor data onto the pial and white matter cortical surfaces and then compares the fit of the two models (Troebinger et al., 2014a). This provides an overall measure of model fit, but cannot provide spatially-specific comparisons. To address this limitation, the ROI analysis reconstructs the data onto both pial and white matter surfaces simultaneously, computes an ROI based on the change of activity on either surface from a baseline time window, and compares the reconstructed activity within the ROI between the two surfaces. We tested four different commonly used functional priors: minimum norm (IID; Hämäläinen and Ilmoniemi, 1984, 1994), LORETA (COH; Pascual-Marqui, 1999; Pascual-Marqui et al., 1994), empirical Bayes beamformer (EBB; Belardinelli et al., 2012; López et al., 2014), and multiple sparse priors (MSP; Friston et al., 2008). These functional priors each embody different assumptions about the distribution of current flow across the cortex, from complete independence (IID), to locally coherent and distributed (COH), to uncorrelated in time (EBB), to locally coherent and sparse (MSP).

## 2 Methods

### 2.1 MRI Acquisition

Data were acquired from a single volunteer with a 3T whole body MR system (Magnetom TIM Trio, Siemens Healthcare, Erlangen, Germany) using the body coil for radio-frequency (RF) transmission and a standard 32-channel RF head coil for reception. A quantitative multiple parameter map (MPM) protocol, consisting of 3 differentially-weighted, RF and gradient spoiled, multi-echo 3D fast low angle shot (FLASH) acquisitions and 2 additional calibration sequences to correct for inhomogeneities in the RF transmit field (Callaghan et al., 2015; Lutti et al., 2012, 2010), was acquired with whole-brain coverage at 800 µm isotropic resolution.

The FLASH acquisitions had predominantly proton density (PD), T1 or MT weighting. The flip angle was 6° for the PD- and MT-weighted volumes and 21° for the T1 weighted acquisition. MT-weighting was achieved through the application of a Gaussian RF pulse 2 kHz off resonance with 4 ms duration and a nominal flip angle of 220° prior to each excitation. The field of view was 256mm head-foot, 224 mm anterior-posterior (AP), and 179 mm right-left (RL). Gradient echoes were acquired with alternating readout gradient polarity at eight equidistant echo times ranging from 2.34 to 18.44 ms in steps of 2.30 ms using a readout bandwidth of 488 Hz/pixel. Only six echoes were acquired for the MT-weighted acquisition in order to maintain a repetition time (TR) of 25 ms for all FLASH volumes. To accelerate the data acquisition, partially parallel imaging using the GRAPPA algorithm was employed with a speed-up factor of 2 in each phase-encoded direction (AP and RL) with forty integrated reference lines.

To maximise the accuracy of the measurements, inhomogeneity in the transmit field was mapped by acquiring spin echoes and stimulated echoes across a range of nominal flip angles following the approach described in Lutti et al. (2010), including correcting for geometric distortions of the EPI data due to B0 field inhomogeneity. Total acquisition time for all MRI scans was less than 30 min.

Quantitative maps of proton density (PD), longitudinal relaxation rate (R1 = 1/T1), magnetisation transfer saturation (MT) and effective transverse relaxation rate (R2* = 1/T2*) were subsequently calculated according to the procedure described in Weiskopf et al. (2013).

### 2.2 FreeSurfer Surface Extraction

FreeSurfer (v5.3.0; Fischl, 2012) was used to extract cortical surfaces from the multi-parameter maps. Use of multi-parameter maps as input to FreeSurfer can lead to localized tissue segmentation failures due to boundaries between the pial surface, dura matter and CSF showing different contrast compared to that assumed within FreeSurfer algorithms (Lutti et al., 2014). Therefore, an in-house FreeSurfer surface reconstruction procedure was used to overcome these issues, using the PD and T1 volumes as inputs. Detailed methods for cortical surface reconstruction can be found in Carey et al. (2017). This process yields surface extractions for the pial surface (the most superficial layer of the cortex adjacent to the cerebro-spinal fluid, CSF), and the white/grey matter boundary (the deepest cortical layer). Each of these surfaces is downsampled by a factor of 10, resulting in two meshes comprising 33,596 vertices each. For the purpose of this paper, we will use these two surfaces to represent deep (white/grey interface) and superficial (grey-CSF interface) cortical models. Cortical thickness was computed as the distance between linked vertices on the pial and white matter surfaces (Kabani et al., 2001; Lerch and Evans, 2005a; MacDonald et al., 2000), and smoothed over each surface with a Gaussian kernel (FHWM=8mm). Mean surface curvature, a measure of local cortical folding, was computed as the mean of the two principal curvatures at each vertex (Davatzikos and Bryan, 1996; Griffin, 1994; Joshi et al., 1995; Luders et al., 2006; Van Essen and Drury, 1997). Sulcal depth was computed using the CAT12 toolbox <u>(http://dbm.neuro.uni-jena.de/cat/)</u> to generate a convex hull surface from the pial surface, and then computing the Euclidean distance between each vertex and the nearest vertex on hull surface (Im et al., 2006; Tosun et al., 2015; Van Essen, 2005).

### 2.3 Simulations

We tested the efficacy of each analysis method using synthetic data sets. All simulations were based on a single dataset acquired from a real experimental recording of the same human participant that the MRI scans were acquired from using a CTF 275 channel Omega system. The MEG sensor data from this dataset were discarded; the dataset was only used to determine the sensor layout, sampling rate (1200Hz, downsampled to 250Hz), number of trials (515), and number of samples (1251) for the simulations. All simulations and analyses were implemented using the SPM12 software package (<u>http://www.fil.ion.ucl.ac.uk/spm/software/spm12/</u>) and are available at <u>http://github.com/jbonaiuto/laminar_sim</u>.

In each simulation, we specified a simulated source centered at a vertex on either the pial or white matter surface. We simulated sinusoidal activity profiles of 20Hz with a patch size of FWHM=5mm over a time window from 100 to 500ms, and used a single shell forward model (Nolte, 2003) to generate a synthetic dataset from the simulated activity. We chose 60 random vertices on each surface to simulate sources at, giving a total of 120 synthetic datasets (60 sources simulated on the pial surface, and 60 on the white matter surface; Figure 1). Each simulation consisted of a single dipole with a moment of 10nAm unless otherwise specified. Typical per-trial SNR levels for MEG data range from -40 to -20dB (Goldenholz et al., 2009), and therefore Gaussian white noise was added to the simulated data and scaled in order to yield per-trial amplitude SNR levels (averaged over all sensors), of -100, -50, -20, -5, -0 or 5dB in order to generate synthetic datasets across a range of realistic SNRs.

**Figure 1:**
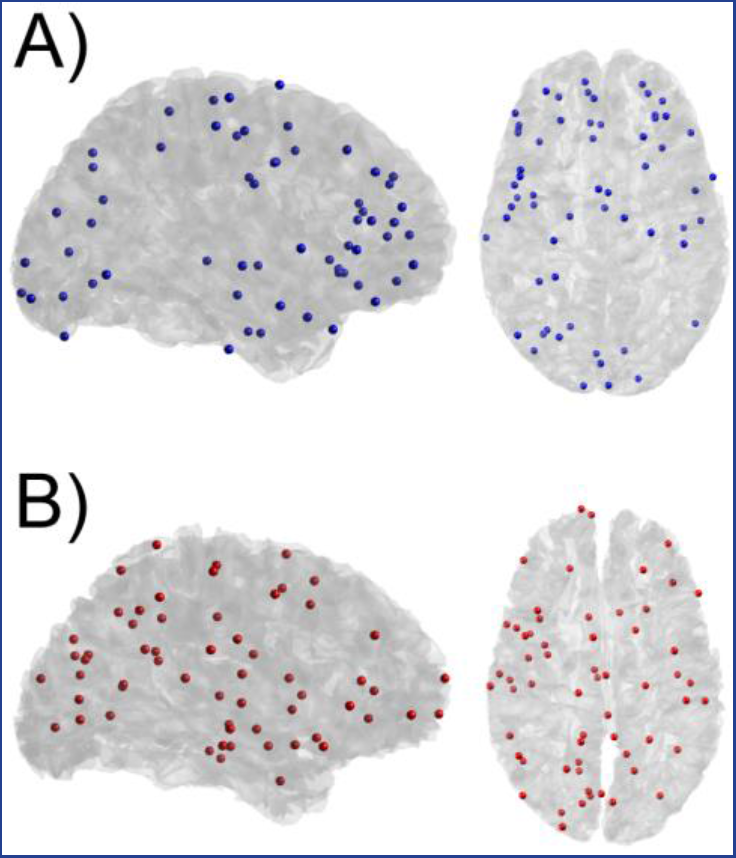
Simulation locations. Locations of simulated activity on the pial surface (A) shown in blue, and the white matter surface (B) shown in red. The vertices were chosen randomly, 60 on the pial surface, and 60 on the white matter surface. All analyses were performed on simulated data from the same 120 source locations.

### 2.4 Analyses for Laminar Discrimination

We compared two methods for determining the laminar locus of simulated activity: a whole brain and a region of interest (ROI) analysis (Figure 2). Each analysis computed 4 different models to explain the simulated data (IID, COH, EBB, and MSP), each using different functional priors expressing common MEG inversion assumptions.

**Figure 2:**
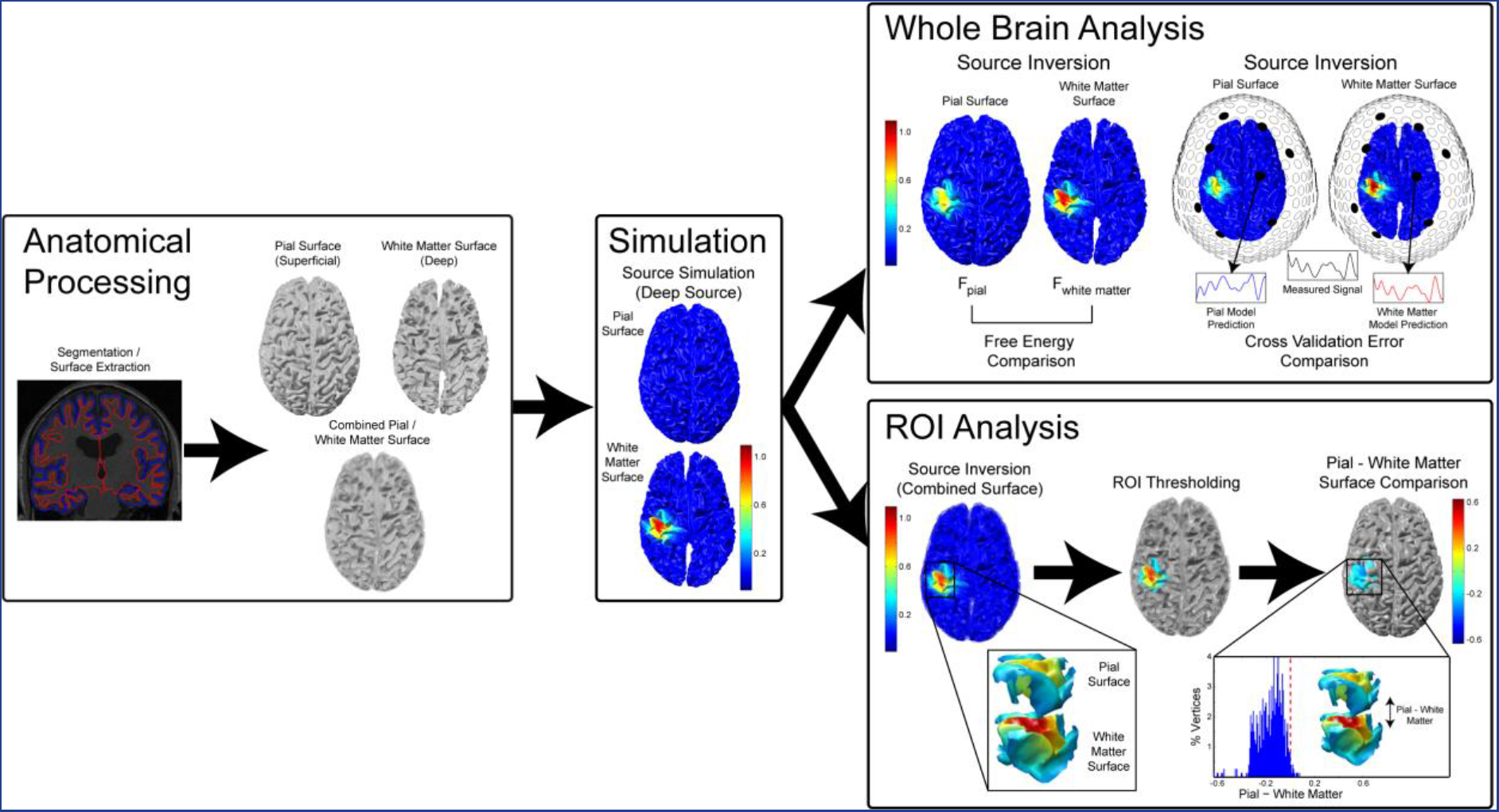
Simulation and analysis protocol. Pial and white matter surfaces are constructed based on an MPM volume. Sources are simulated as patches of 20Hz activity on random vertices of either the pial or white matter surface. The whole brain analysis computes separate generative models for each surface and compares these models in terms of free energy or cross validation error. The ROI analysis creates a single generative model combining both surfaces. It then creates an ROI localization of baseline-corrected increase in power, and compares estimated power at corresponding vertices between the surfaces within that ROI.

The whole brain analysis reconstructed the simulated data (with sensor noise) separately onto each of the surface models (the pial and the white matter) and compared the fit between the two models using either free energy (Troebinger et al., 2014a) or cross validation error. Free energy is a parametric metric that rewards fit accuracy and penalizes model complexity (Friston et al., 2007), providing a lower bound for the log model evidence value (Penny et al., 2010). Cross validation involves partitioning the data into training and test portions. The idea is to fit models to the training data and then to compare models based on their accuracy in predicting the test data. Models which are too complex will over-fit the training data (i.e. fit the noise) and therefore perform poorly on the test data. Conversely, models which are too simple will not be able to explain the training or the test data. In other words, cross validation involves a similar accuracy-complexity trade-off to free energy, but is calculated on the basis of the sensor-level time-courses, rather than the distance between the means of the prior and posterior distributions. This in turn means that while free energy is dependent on the prior distribution specified (including the variance thereof), cross validation is not. We computed the average 10-fold cross validation error by excluding 10% of the sensors from the source reconstruction and computing the error in predicting the missing sensor data using the resulting model. The error in each fold was defined as the root mean square error (RMSE) of the sensor data predictions, expressed as a percentage of the root mean square (RMS) measured sensor data, averaged over each excluded sensor. Much like arguments for parametric and non-parametric statistics, the free energy approximation is more powerful (as it uses all the data) when the underlying assumptions are met, whereas cross validation is not quite as sensitive, is more time consuming, yet is robust.

The ROI analysis reconstructed the data (with sensor noise) onto a mesh combining the pial and white matter surfaces, thus providing an estimate of source activity on both surfaces. We defined an ROI by comparing power in the 10-30Hz frequency band during the time period containing the simulated activity (100ms to 500ms) with a prior baseline period (−500ms to −100ms) at each vertex using paired t-tests. Vertices in either surface with a t-statistic in the 75^th^ percentile of the t-statistics over all vertices in that surface, as well as the corresponding vertices in the other surface, were included in the ROI. This ensured that the contrast used to define the ROI was orthogonal to the subsequent pial versus white matter surface contrast. For each trial, ROI values for the pial and white matter surfaces were computed by averaging the absolute value of the change in power compared to baseline in that surface within the ROI. Finally, a paired t-test was used to compare the ROI values from the pial surface with those from the white matter surface over trials. All t-tests were performed with corrected noise variance estimates in order to attenuate artifactually high significance values (Ridgway et al., 2012).

Two-sided binomial tests were used to compare the accuracy in classifying simulated sources as originating from the correct surface, as well as bias toward the pial surface, with chance levels (50%). The whole brain and ROI analyses as well as source reconstruction algorithms were compared in terms of their classification accuracy using exact McNemar’s tests (McNemar, 1947). Correlations between free energy and cross validation error differences, and ROI t-statistics were evaluated using Spearman’s rho tests. The free energy difference was computed as the free energy for the pial surface model minus that of the white matter surface model, while the cross validation error difference was computed as the cross validation error for the white matter surface model minus that of the pial surface model. This ensured that for both metrics, positive values indicate a better fit metric for the pial surface model. Relationships between the difference in free energy between the correct and incorrect surface models, and cortical surface statistics such as cortical thickness, mean surface curvature, sulcal depth, and lead field strength were evaluated using Spearman’s rho tests. Because the surface statistics were all potentially correlated with each other, the correlation coefficients were compared using Meng’s test for correlated correlation coefficients (Meng et al., 1992), followed up by pairwise Z-tests.

### 2.5 Source reconstruction

Source inversion was performed using four different algorithms within SPM12: empirical Bayesian beamformer (EBB; Belardinelli et al., 2012; López et al., 2014), minimum norm (IID; Hämäläinen and Ilmoniemi, 1984, 1994), LORETA (COH; Pascual-Marqui, 2002; Pascual-Marqui et al., 1994) and multiple sparse priors (MSP; Friston et al., 2008). The source inversion in the whole brain analysis was applied to a Hann windowed time window from −500ms to +500ms filtered to 10-30Hz, while the ROI analysis did not use a Hann window. These data were projected into 274 orthogonal spatial (lead field) modes and 4 temporal modes.

The empirical Bayes optimization rests upon estimating hyper-parameters which express the relative contribution of source and sensor level covariance priors to the data (López et al., 2014). For all algorithms we assumed the sensor level covariance to be an identity matrix. For the EBB and COH algorithms there is a single source level prior which is either estimated from the data (EBB) or fixed (COH). There are therefore only two hyper-parameters to estimate – defining the relative contribution of the source and sensor level covariance components to the data. For the MSP algorithm, there is a different source covariance prior for every possible patch. We used a total of 90 patches, 60 of which were used as locations for potential simulated sources, plus 30 patches at random vertices. This does give MSP a considerable advantage (see Discussion) but factors out computational issues in the optimization.

## 3 Results

### 3.1 Laminar source discrimination

In the whole brain analysis, we computed a difference in free energy between the pial and white matter generative models, approximating the log ratio of the model likelihoods. This resulted in a metric which is positive or negative if there is more evidence for the pial or white matter model, respectively. Similarly, the ROI analysis produced a t-statistic which was positive when the change in power was greater on the pial surface, and negative when the change was greater on the white matter surface. The free energy difference and ROI t-statistics for each simulation using each source inversion algorithm are shown in Figure 3. For the EBB and MSP algorithms (Figure 3A, D), most of the sources simulated on the pial surface resulted in positive free energy differences and t-statistics, while most of those simulated on the white matter surface yielded negative metrics (EBB whole brain: accuracy=93.33%, *p*<0.0001; EBB ROI: accuracy=99.17%, *p*<0.0001; MSP whole brain: accuracy=100%, *p*<0.0001; MSP ROI: accuracy=100%, *p*<0.0001). In contrast, the COH and IID algorithms were biased deep for free energy (COH: white matter=69.17%, *p*<0.0001; IID: white matter=67.5%, *p*<0.0005) and superficial for the ROI analysis (COH: pial=100%, *p*<0.0001; IID: pial=100%, *p*<0.0001) regardless of the layer on which sources were simulated. Thus, both the EBB and MSP versions of both the whole brain and ROI analyses were able to distinguish between white matter and pial sources, but the COH and IID algorithms could not (Figure 3B, C).

**Figure 3:**
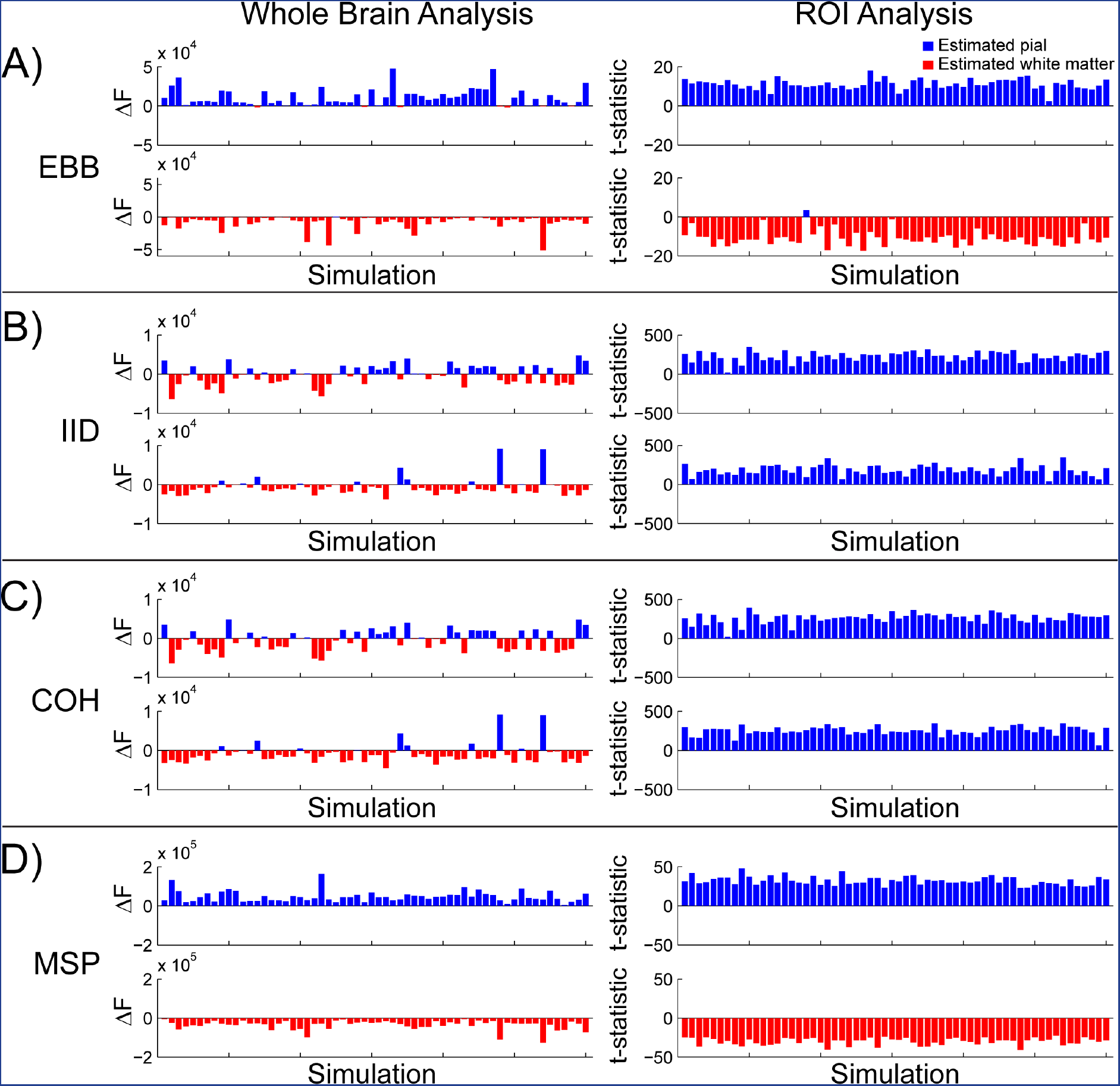
Laminar source discrimination. Left column: The difference in free energy between the pial and white matter generative models in each simulation (SNR=−20dB). Right column: T-statistics from the ROI analysis comparing pial and white matter ROIs for each simulation (SNR=−20dB) Each panel shows simulations with pial surface sources on the top row, and simulations with white matter surface sources on the bottom row. In both the whole brain and ROI analyses, the EBB (A) and MSP (D) source reconstruction algorithms were able to correctly classify simulated source activity as arising from either the deep or superficial surface (EBB whole brain: accuracy=93.33%, p<0.0001; EBB ROI: accuracy=99.17%, p<0.0001; MSP whole brain: accuracy=100%, p<0.0001; MSP ROI: accuracy=100%, p<0.0001). The IID (B) and COH (C) versions of the whole brain analysis were biased toward the deep surface (IID: white matter=67.5%; p<0.0005; COH: white matter=69.17%, p<0.0001), and toward the superficial surface in the ROI analysis (IID: pial=100%, p<0.0001; COH: pial=100%, p<0.0001).

In order to quantify the agreement between the different metrics used (free energy, cross validation error, and ROI t-statistic), we analyzed pairwise correlations between them. The difference in free energy from the whole brain analysis and the t-statistic from the ROI analysis were correlated for the EBB algorithm (ρ(118)=0.75, *p*<0.001; Figure 4A). There was a considerable separation between the MSP t-statistic distributions for pial and white matter sources, so we therefore considered the pial and white matter simulation sources separately. Correlations between the free energy difference and ROI analysis t-statistic were still significant for sources on both surfaces (pial: ρ(58)=0.48, *p*<0.0005, white matter: ρ(58)=0.3, *p*=0.02; Figure 4B). We compared the difference in cross validation error between the white matter and pial surface models with the difference in free energy in order to verify that the results of the whole brain analysis were the same with an independent metric. We found that the cross validation error difference was highly correlated with both the free energy difference (EBB: ρ(118)=1.0, *p*<0.0001; Figure 4A; MSP: ρ(118)=0.98, *p*<0.0001; Figure 4B) and the t-statistic from the EBB version of the ROI analysis (ρ(118)=0.75, *p*<0.0001; Figure 4A). The cross validation error difference and ROI t-statistic were only significantly correlated for pial sources for the MSP algorithm (pial: ρ(58)=0.34, *p*=0.009, white matter: ρ(58)=0.18, *p*=0.173; Figure 4B), but the two metrics predicted the same classification category (pial or white matter) for every simulated source.

**Figure 4:**
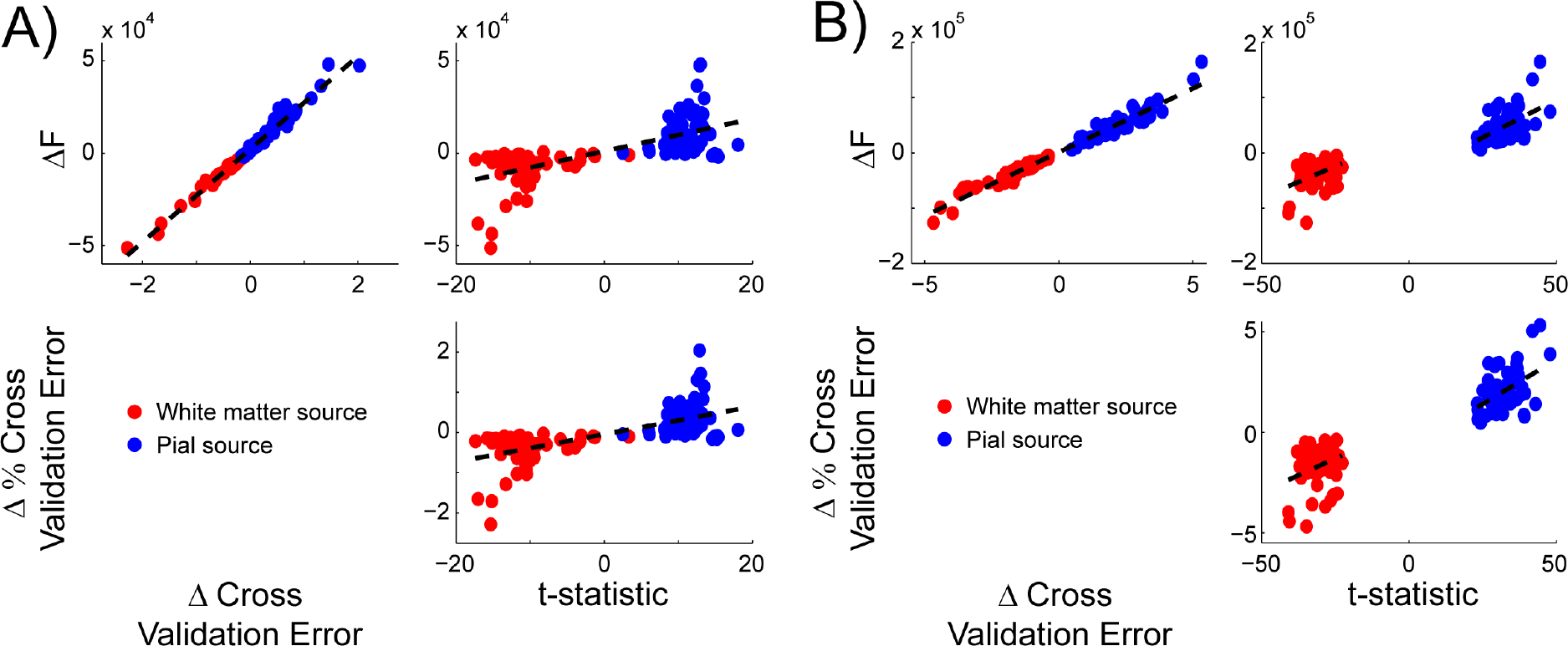
Metric correlations. All of the analysis metrics (free energy difference, cross validation error difference, and ROI t-statistic) were highly correlated with each other for both the EBB (A) and MSP (B) source inversion algorithms (SNR=−20dB). Red circles denote sources simulated on the white matter surface, blue circles denote those simulated on the pial surface. Dotted lines indicate the least squares fitted linear relationships. The separated dashed lines in panel B indicate that the metric correlations were performed within (not across) layers in these cases.

At an SNR of −20dB, the EBB and MSP versions of the whole brain and ROI analyses correctly classified most of the simulated sources, while the IID and COH versions performed at chance levels (Figure 3). In order to further evaluate the performance of the analyses, we simulated sinusoidal activity with varying levels of noise resulting in SNRs from −100dB to 5dB, as well as a control dataset containing only noise and no signal (SNR=-∞dB). We first compared the bias of each analysis and source inversion algorithm by computing the percentage of sources classified as pial, using a threshold of ±3 for the free energy difference (meaning that one model is approximately twenty times more likely than the other) and a threshold of the critical t value with *df*=514 and α=0.05 for the ROI t-statistic. At low levels of SNR, all of the metrics were biased toward pial sources (SNR=−100dB, EBB whole brain: pial=85%, *p*<0.0001; SNR=−100dB, EBB ROI: pial=100%, *p*<0.0001; SNR=−100dB, MSP whole brain: pial=74.17%, *p*<0.0001), except for the MSP version of the ROI analysis (SNR=−100dB: pial=42.5%, *p*=0.12), but these biases were not statistically significant (i.e. the free energy difference and t-statistics were greater than zero, but did not exceed the significance threshold). As SNR increased nearly all classifications exceeded the significance threshold, and the EBB and MSP versions of the whole brain and ROI analyses became unbiased (SNR=5dB, EBB whole brain: pial=50%, *p*=1.0; SNR=5dB, EBB ROI: pial=50%, *p*=1.0; SNR=5dB, MSP whole brain: pial=50%, *p*=1.0; SNR=5dB, MSP ROI: pial=50%, *p*=1.0; Figure 5A, C). We then compared the accuracy of each analysis and source inversion algorithms over the range of SNRs.

**Figure 5:**
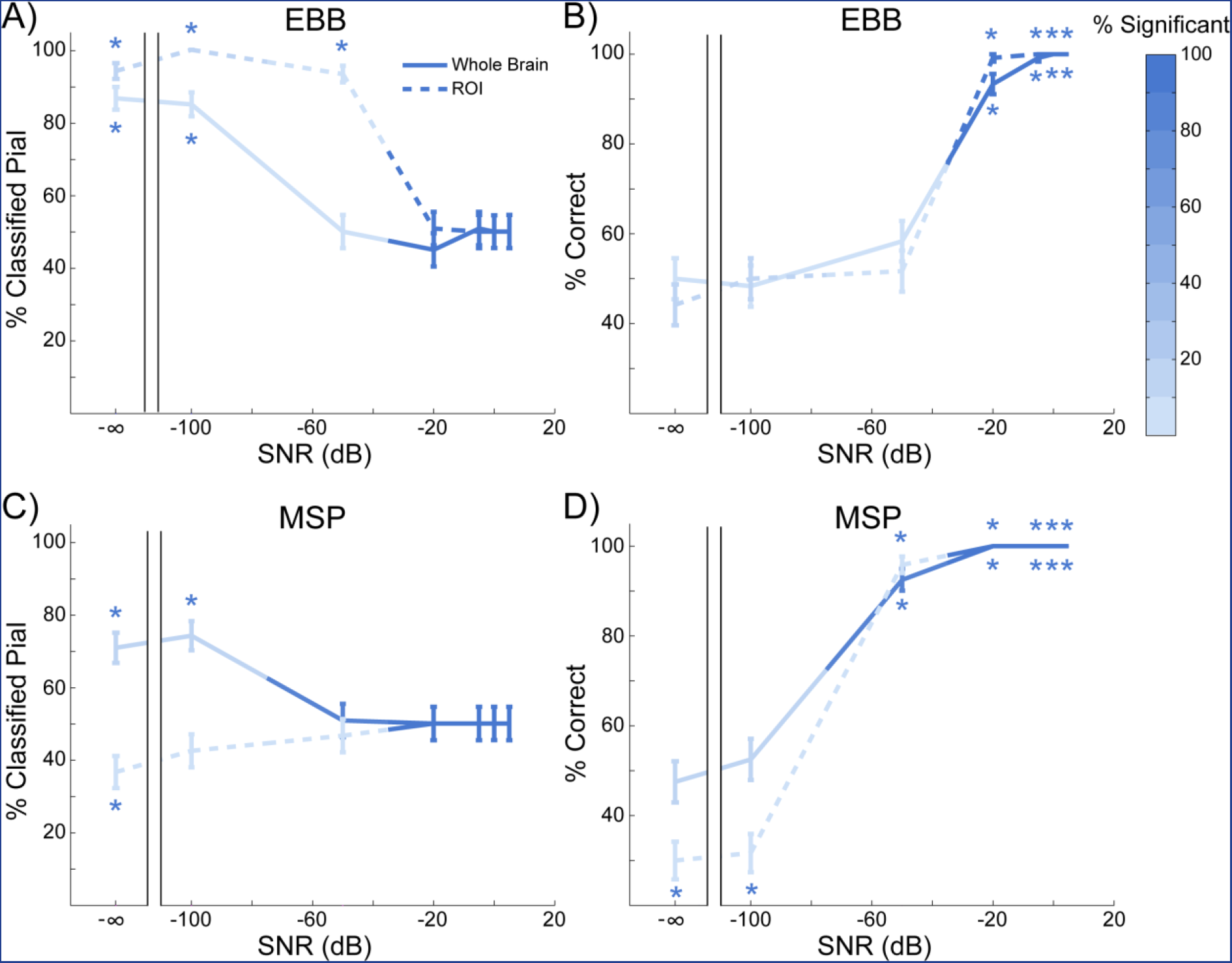
Laminar discrimination accuracy. A) The percentage of sources classified as originating from the pial surface for the EBB version of the whole brain and ROI analyses, for each level of SNR tested. The error bars represent the standard error. The percentage of simulations with free energy differences or t-statistics exceeding the significance threshold is represented by the intensity of the line color. Asterisks show where the percentage is significantly above or below chance levels. Both analyses were biased toward the pial surface at low SNR levels, but not significantly. As SNR increased, nearly all of the classifications exceeded the significance threshold and both analyses became unbiased. B) The percentage of sources accurately classified by the EBB version of the whole brain and ROI analyses, over all tested SNR levels. Both analyses accurately classified at least 90% of the simulated sources at SNR=−20dB. C) and D) As in (A) and (B) for the MSP version of both analyses. The MSP version of the ROI analysis was unbiased even at low SNR levels and the whole brain analysis correctly classified at least 90% of the simulated sources at SNR=−50dB.

The MSP version of the whole brain analysis performed at above 90% accuracy (thresholded) even with SNR=−50dB (accuracy=92.5%, *p*<0.0001; Figure 5D), while the MSP version of the ROI analysis and the EBB versions of both analyses required at least −20dB SNR to achieve approximately 90% accuracy (MSP ROI: accuracy=100%, *p*<0.0001; EBB ROI: accuracy=99.17%, *p*<0.0001; EBB whole brain: 93.33%, *p*<0.0001; Figure 5B, D). The MSP version of the whole brain analysis outperformed the ROI analysis at SNR=−50dB (*Χ*^2^(1, N=120)=97.01, *p*<0.0001), and the EBB version of the whole brain analysis at SNR=−50dB (*Χ*^2^(1, N=120)=96.01, *p*<0.0001) and SNR=−20dB (*Χ*^2^(1, N=120)=6.13, *p*=0.013). All differences between the EBB and MSP algorithms and analyses disappeared at higher SNR levels.

Because we scaled the level of the white noise in order to achieve a fixed SNR level, there was the possibility that these results were biased due to boosting the signal from deep layer sources which were further away from the sensors. In order to address this, we ran additional control simulations in which we kept the source signal magnitude fixed at 20nAm, and varied the level of the white noise from 10 to 2×10^6^ fT RMS. Consistent with the results of our main simulations, both the EBB and MSP versions of the whole brain and ROI analyses achieved at least 80% accuracy when the noise level fell below 100 fT RMS (EBB whole brain: 80.83%, *p*<0.0001; EBB ROI: 90.83%, *p*<0.0001; MSP whole brain: 99.17%, *p*<0.0001; MSP ROI: 100%, *p*<0.0001; **Figure S1**). For comparison, this corresponds to a per-trial SNR level of approximately −25 dB.

We tested the EBB and MSP versions of the whole brain and ROI analyses under more realistic noise assumptions by simulating sources added to resting state MEG data acquired from the same participant whose anatomy the simulations were based on. Data from a 10 minute resting state scan with the eyes open were downsampled to 250Hz, epoched to match the simulated trial durations, and baseline corrected to remove the DC offset. We varied the strength of the simulated dipoles from 0.01nAm to 300nAm (resulting in a range of SNRs from approximately −100dB to −5dB), positioned them at the same vertices as the main simulations, and added their projected sensor activity to the resting state data (**Figure S2**). The EBB and MSP versions of the whole brain and ROI analyses achieved significantly above chance accuracy when the dipole moment was at least 50nAm (corresponding to an SNR of approximately −20 dB), MSP was more accurate than EBB, and the ROI analysis was more accurate than the whole brain analysis, consistent with the main simulation results (**Figure S3**). Note, however, that at low dipole moment both the EBB and MSP global metrics are deemed significant yet are biased towards the superficial surface (see Discussion).

### 3.2 Patch size

We modelled current flow as normal to the cortical surface, but the spatial extent (or local dispersion) of this current flow tangential to the cortical surface is an uncertain quantity which will depend on a number of factors including lateral local connectivity. In our previous work we have shown that incorrect estimates of cortical patch size tend to bias layer estimates (Troebinger et al., 2014a). We therefore simulated dispersions (or patch sizes) of current flow over the cortical surface of 5mm and 10mm, and tested the whole brain analysis using source reconstruction patch sizes of 5mm and 10mm. As SNR increased the EBB and MSP versions of the whole brain analysis went from being biased to the pial surface (SNR=-∞dB, simulate=5mm, reconstruct=5mm, EBB: pial=85%, *p*<0.0001; SNR=-∞dB, simulate=10mm, reconstruct=10mm, EBB: pial=85%, *p*<0.0001; SNR=-∞dB, simulate=5mm, reconstruct=5mm, MSP: pial=74.17%, *p*<0.0001; SNR=-∞dB, simulate=10mm, reconstruct=10mm, MSP: pial=77.5%, *p*<0.0001) to being unbiased when the patch size was correctly estimated (SNR=5dB, simulate=5mm, reconstruct=5mm, EBB: pial=50%, *p*=1.0; SNR=5dB, simulate=10mm, reconstruct=10mm, EBB: pial=51.67%, *p*=0.784; SNR=5dB, simulate=5mm, reconstruct=5mm, MSP: pial=50%, *p*=1.0; SNR=5dB, simulate=10mm, reconstruct=10mm, MSP: pial=50%, *p*=1.0; Figure 6A, C). However, when the patch size was either under- or over-estimated, the EBB algorithm became biased toward the white matter surface (SNR=5dB, simulate=5mm, reconstruct=10mm: pial=32.5%, *p*<0.001; SNR=5dB, simulate=10mm, reconstruct=5mm: pial=32.5%, *p*<0.001), while the MSP algorithm remained unbiased (SNR=5dB, simulate=5mm, reconstruct=10mm: pial=55%, *p*=0.315; SNR=5dB, simulate=10mm, reconstruct=5mm: pial=55%, *p*=0.315). As a result, the laminar classification accuracy of both algorithms was reduced when the patch size was either under- (SNR=−50dB, MSP: *Χ*^2^(1, N=120)=6.05, *p*=0.014; SNR=−20dB, EBB: *Χ*^2^(1, N=120)=17.05, *p*<0.0001) or over-estimated (SNR=−50dB, MSP: *Χ*^2^(1, N=120)=6.86, *p*=0.009; SNR=−20dB, EBB: *Χ*^2^(1, N=120)=33.03, *p*<0.0001; Figure 6B, D), although MSP was less sensitive to this difference. The accuracy of the EBB and MSP versions of the ROI analysis were similarly affected by the patch size estimate (**Figure S4**).

**Figure 6:**
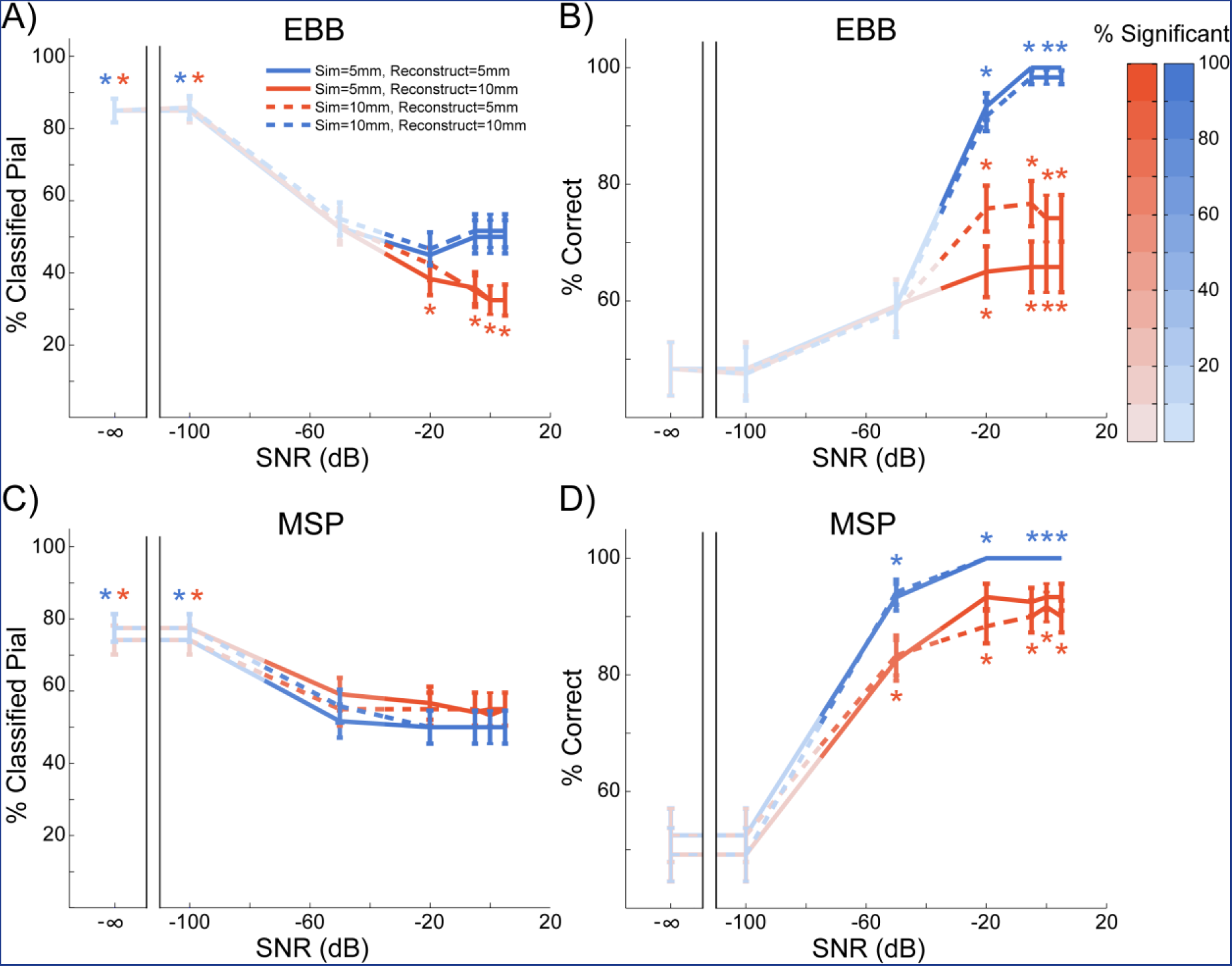
Whole brain analysis performance with varying patch sizes. Blue lines denote simulations where the reconstructed patch size matches the simulated patch size (solid=5mm, dashed=10mm), red lines are where patch size is either under- (dashed red) or over-estimated (solid red). The percentage of sources significantly classified as either pial or white matter is represented by the intensity of the line color. Asterisks show where the percentage is significantly above or below chance levels. A) The percentage of sources classified as originating from the pial surface for the EBB version of the whole brain analysis, for each level of SNR tested. The error bars represent the standard error. As SNR increased, simulations where the patch size was correctly estimated became unbiased, while incorrect patch size estimates resulted in a bias toward the white matter surface. B) Incorrect patch size estimates resulted in reduced classification accuracy for the EBB version of the whole brain analysis. C) The MSP version of the whole brain analysis became biased toward the pial surface as SNR increased. D) Classification accuracy was reduced for the MSP version of the whole brain analysis when the patch size was under- or over-estimated.

### 3.3 Surface anatomy

We next sought to determine what anatomical features of the cortical surface make it easier or more difficult to discriminate between white matter and pial surface sources. We therefore computed several surface statistics including cortical thickness (Kabani et al., 2001; Lerch and Evans, 2005a; MacDonald et al., 2000), surface mean curvature (Davatzikos and Bryan, 1996; Griffin, 1994; Joshi et al., 1995; Luders et al., 2006; Van Essen and Drury, 1997), sulcal depth (Im et al., 2006; Tosun et al., 2015; Van Essen, 2005), and lead field RMS (Hillebrand and Barnes, 2002), and examined the relationship between each measure and the difference in free energy between the correct and incorrect generative models in the whole brain analysis (Δ*F* = *F*_*correct*_ − *F*_*incorrect*_, Figure 7). The distributions of cortical thickness, mean curvature, and sulcal depth closely matched previously published estimates (Fischl and Dale, 2000; Hutton et al., 2008; Jones et al., 2000; MacDonald et al., 2000; Tosun et al., 2015). Sulcal depth and lead field RMS were both highly correlated with Δ*F* (sulcal depth: ρ(118)=−0.726, *p*<0.0001; lead field RMS: ρ(118)=0.653, *p*<0.0001), indicating that source activity was more easily classified as originating from the pial or white matter surface in sources closer to the scalp, and those which have a greater impact on the sensors. Cortical thickness and surface curvature were both weakly correlated with Δ*F* (cortical thickness: ρ(118)=0.288, *p*<0.005; surface curvature: ρ(118)=0.294, *p*<0.005), meaning that source activity was more easily classified as pial or white matter where there was greater distance between the surfaces, and at gyral rather than sulcal vertices. These surface metrics are not independent (e.g. both surface curvature and sulcal depth influence the lead field RMS), but a comparison of correlated correlation coefficients (Meng et al., 1992) revealed significant heterogeneity of the absolute correlation matrix (*Χ*^2^(3, N=120)=48.72, *p*<0.0001). Follow-up tests revealed that sulcal depth and lead field RMS were significantly more correlated with Δ*F* than either cortical thickness (sulcal depth: Z=5.6, *p*<0.0001; lead field RMS: Z=4.37, *p*<0.0001) or surface curvature (sulcal depth: Z=5.34, *p*<0.0001; lead field RMS: Z=4.12, *p*<0.0001) were, and there was no difference between the sulcal depth and lead field RMS correlation coefficients (Z=1.22, *p*=0.111).

**Figure 7:**
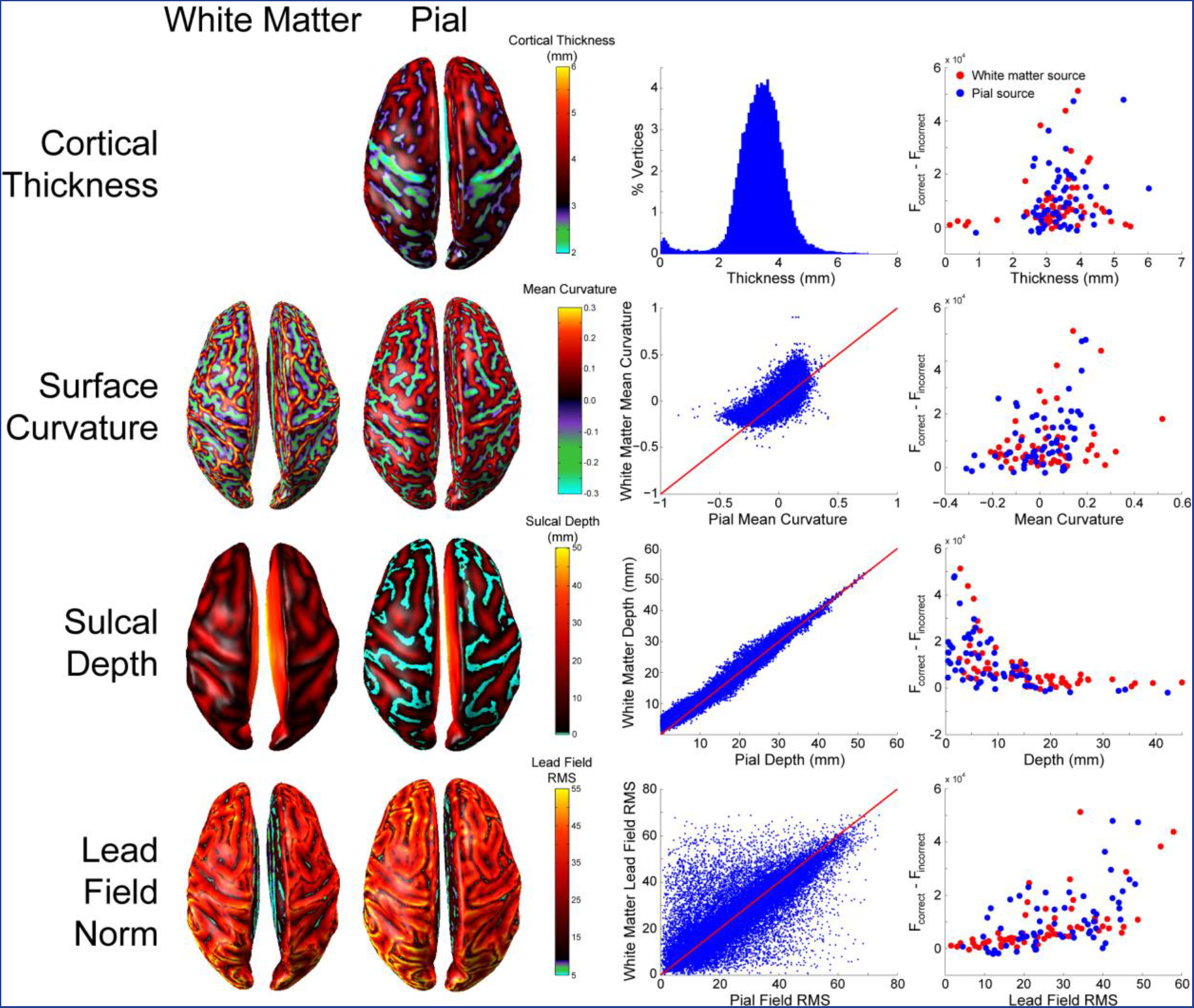
Relationship between cortical surface anatomy and laminar discriminability. First row (from left to right): cortical thickness plotted on the pial surface, cortical thickness distribution over the pial surface, free energy difference between the correct and incorrect generative model as a function of cortical thickness (red = white matter sources, blue = pial sources; EBB, SNR=−20dB). Second row: Surface curvature of the white matter and pial surfaces, relationship between white matter and pial surface curvature, free energy difference as a function of surface curvature. Third row: Sulcal depth of the white matter and pial surfaces, relationship between white matter and pial surface sulcal depth, free energy difference as a function of sulcal depth. Fourth row: Lead field RMS of the white matter and pial surfaces, relationship between lead field RMS of the white matter and pial surfaces, free energy difference as a function of lead field RMS.

We also looked at the distribution of lead-field differences between matching pial/white vertices relative to their nearest neighbours on the same surface (**Figure S8**) and found that generally (in 66.8% on the pial surface and 56.65% on the white matter surface) of vertices there was a smaller difference in lead-field magnitude between pial-white vertex pairs than their immediate same-surface neighbours.

## 4 Discussion

We here provide a comparison and evaluation of analysis techniques for non-invasive laminar specific inference in human neocortex with MEG. We found that, given sufficient SNR, both the whole brain model comparison analysis and the ROI analysis were able to distinguish between simulated activity originating on white matter versus pial surfaces, representing deep and superficial cortical laminae, from the sensor data alone. Importantly, we found mutually corroborating results from two inversion schemes (MSP and EBB) and three metrics of fit (free energy, cross-validation and local t-tests). The MSP source inversion algorithm was more sensitive to laminar differences in activity at a lower SNR, though EBB achieved similar classification performance with a slightly higher SNR. These SNR levels are now possible thanks to novel head-cast technology which reduces head movement and allows accurate co-registration over repeated sessions, resulting in very high SNR MEG data (Meyer et al., 2017; Troebinger et al., 2014a, 2014b).

### 4.1 Laminar Discrimination Depends on Functional Assumptions

In this study we remained agnostic about functional assumptions and used four commonly used sets of priors. We find that only the algorithms with some sparsity constraint (EBB and MSP) could successfully discriminate the laminar origin of source activity (Figure 3). This is because model evidence, like cross-validation, penalizes more complex models as they tend to have poor generalization performance. This means that intrinsically sparser algorithms (like EBB and MSP) will always be rewarded if they can explain the same amount of data with fewer active sources. We should note that MSP had a distinct advantage here as the possible source space of priors included the 120 vertex locations on which sources were simulated. EBB had no such prior information. We found that the implementations of minimum norm and LORETA were not suitable for this discrimination task, but we should point out that these algorithms (as implemented in SPM12) are somewhat generic and not individually optimized. For example, many groups define a baseline period or empty room recording, which allows an estimate of the optimal regularization parameter, or use a depth re-weighting in the minimum norm estimates (Gramfort et al., 2014). Here all regularization (the balance between the source and sensor level covariance matrices) was set based on a Free energy optimization (Friston et al., 2008), but cross-validation approaches are also possible (Engemann and Gramfort, 2015).

We note that the ROI and global metrics implemented here are not based on identical data. The ROI analysis requires a functional contrast – it can only detect laminar differences where there is a change in power from a baseline time window, while the whole brain analysis simply determines which surface best supports the measured data (even if there is no modulation from baseline). Another difference in the implementation is the use of a Hann window in the whole brain analyses (originally motivated to give frequency specificity) which was not present in the local analysis and hence the local metrics have a marginal SNR advantage.

### 4.2 Laminar Discrimination Requires Accurate Patch Size Estimates

We looked at a range of simulation and reconstruction patch sizes, as previous simulation work has shown that an overestimation of patch extent can bias model evidence towards superficial cortical layer models (Troebinger et al., 2014a). In this study we observed a similar skew in the mean free energy difference (**Figure S5**) but this was always much smaller than the free energy difference for the correct surface. This is perhaps due to the refined and much smoother surfaces we are using here (and that we are assuming zero co-registration error). Here we focused on accuracy (rather than mean free energy difference) of laminar classification and found that it was indeed degraded when the source patch size was over- or under-estimated, but that EBB (Figure 6B) was biased toward white matter sources, while MSP was slightly biased toward pial sources (Figure 6A,C), regardless of the over- or under-estimation. EBB is a two stage process, first estimating the source distribution with beamformer priors, and then balancing this estimate against sensor noise to fit the data. For a single source, under- or over-estimation of patch size using EBB will lead to a source estimate with a lower peak variance (Hillebrand and Barnes, 2011). In the limit this will tend to a flat variance distribution across the cortex (resembling an IID or COH prior), and our initial simulations (Figure 3) show that free energy metrics of these estimates (at high SNR) will be biased toward the deep layers (**Figure S6**). We speculate that given similar levels of accuracy, free energy favors less the marginally less complex current distributions on the deep layer models. In contrast, laminar classification based on the percentage of variance explained, a goodness of fit metric that does not penalize complexity, results in a superficial bias for IID and COH (**Figure S7**). We also note that the ROI analysis for IID and COH has a superficial bias (Figures 3, S6).

### 4.3 Free Energy and Cross-Validation Error are Closely Related

Free energy is a widely used parametric measure of model fit in the neuroimaging community, but cross validation error is a more commonly used nonparametric measure in the field of machine learning. Both metrics reward model accuracy and try to avoid overfitting of data. Free energy accomplishes this by penalizing model complexity, while cross validation error measures the ability of a model to generalize to new, unseen data. In our simulations, the difference in cross validation error between the pial and white matter generative models was highly correlated with their difference in free energy. Importantly, this demonstrates that the laminar inferences in these analyses are not specific to specialized parametric measures of model fit such as free energy, but generalize to familiar nonparametric measures such as cross validation error.

### 4.4 Anatomical Assumptions

We made three major simplifying anatomical assumptions in these analyses: i) the locations of deep and superficial laminar sources are on the white matter and pial surfaces, ii) deep and superficial laminae contribute equally to the measured MEG signal, and iii) the spatial spread of lateral connectivity is the same across laminae. However, these assumptions may require further scrutiny. First, the mean thickness of the human cerebral cortex ranges from 2mm to over 4mm (Lerch and Evans, 2005b; MacDonald et al., 2000), and therefore the effective net dipole moments of sources in the supra- and infra-granular layers will be more proximal than the pial and white matter surfaces used here. Second, while the main contributors to the MEG signal are the supra-granular layers II/III pyramidal neurons and the infra-granular layer V pyramidal neurons of the neocortex (Murakami and Okada, 2006) with relatively comparable numbers of neurons in these layers (Meyer et al., 2010), based purely on histology, one would expect the layer V cells to have an approximately 3-4 times larger dipole moment (or impact on the MEG sensors) than the layer II/III cells (Bush and Sejnowski, 1993; Jones et al., 2007; Murakami and Okada, 2015, 2006). Based on geometry, as the layer II/III cells are nearer to the MEG sensors, one might expect this to mitigate the effect. However the regions in which the superficial layers are closer to the sensors (the crests of the gyri) are also the regions in which the MEG signal is attenuated due to their predominantly radial orientation (Hillebrand and Barnes, 2002). In practice, one should expect a slight bias towards the superficial surface (on average the lead fields from the superficial layer are marginally (~12%) greater than the deep - see Figure 7) but this is insignificant as compared to the bias against supra-granular cells due to their size (a factor of 2-4). Third, there are differences in the extent of lateral connectivity in different cortical layers (Kritzer and Goldman-Rakic, 1995; Schubert et al., 2007), and regions (Amir et al., 1993; Elston and Rosa, 1997; Elston et al., 1999; Levitt et al., 1993; Lund et al., 1993). Specifically, superficial layer II/III neurons are generally more sparsely and less locally connected than the deep ones (Sakata and Harris, 2009; Schubert et al., 2007).

### 4.5 Limitations

While the simulations reported here demonstrate the feasibility of laminar specific inferences with MEG, there are several limitations that could be addressed in future work.

Here, each simulation involved a single source of activity on either the white matter or pial surface. In the human brain, multiple sources are often co-active (Hari and Salmelin, 1997; Jensen and Vanni, 2002), and laminar recordings from non-human primates suggest that activity occurs simultaneously at deep and superficial sources in the same patch of cortex (Bollimunta et al., 2011; Haegens et al., 2015; Maier et al., 2010; Smith et al., 2013; Spaak et al., 2012; Xing et al., 2012). The ROI analysis could easily be extended to handle multiple sources across the brain by defining multiple ROIs based on clusters of activity. However, in the case of simultaneous activity in deep and superficial laminae within a cortical patch, both the whole brain and ROI analyses implemented here can only infer the laminar origin of the strongest source. One option could be to make use of biophysically informed generative models such as the canonical microcircuit (Bastos et al., 2012), in order to predict the expected time-series from the different pyramidal cell populations. These could then be projected from different spatial origins, and thus would give a complete spatio-temporal model of the MEG signal.

In the main simulations reported here, we used Gaussian white noise scaled in each simulation in order to achieve a fixed per-trial SNR, which could have biased the results by attenuating noise levels in deep layer simulations. In additional control simulations we have shown that the whole brain and ROI analyses can still make accurate laminar inferences when the SNR is not fixed (**Figure S1**), and with more realistic noise sources (**Figure S2, S3**). However, in all of our simulations we assume that the noise covariance matrix is diagonal, but for real data this would need to be estimated using empty room measurements.

We should point out that the SNR of the dataset, although apparently low, will be significantly increased through the singular value decomposition implicit in the source reconstruction. Here the 251 samples were reduced to 4 orthogonal temporal modes giving an effective SNR increase of a factor of 7.922.

In human MEG data, the true source patch size is generally unknown. Our simulations show that spatial specificity can be reduced by inaccurate estimates of source patch size, and therefore any empirical MEG studies of laminar-specific activity should determine the appropriate patch size in order to control for potential biases. This could be accomplished by using the global fit metrics (free energy or cross validation error) to compare source inversions performed on combined pial/white manifolds for multiple patch sizes, prior to any layer comparison. With the patch size optimized one could then proceed with either the global or local metrics as outlined in this paper.

In these simulations we should note that MSP generally performs the best for two reasons. Firstly, the source configurations simulated were sparse, favoring both MSP and EBB. Secondly, in the MSP case the source space was constrained to the 90 possible simulation locations. We did this to factor out the computationally intensive optimization for a global (free energy) maximum as MSP selects and removes patch combinations. In practice, however, we would typically use multiple initial patch selections (e.g. Troebinger et al., 2014a) to produce multiple solutions from which we would select the solution with highest overall free energy.

The simulations reported here do not contain co-registration error or within-session movement. Previous work suggests that the co-registration error must be less than 2mm/2° in order to make accurate laminar inferences (Troebinger et al., 2014a). While subject-specific headcasts have been shown to achieve this level of precision (Meyer et al., 2017), it is not clear what range of error and within-session movement the specific analyses described here can tolerate.

Finally, both the ROI and whole-brain metrics are biased superficially for low SNR data (Figure 5). In our Gaussian noise simulations this is not an issue, as these metrics are non-significant at these levels. However, in real recording situations it will be important to consider whether a superficial estimate of activity is not to mischaracterization of the noise (see **Figure S3**, the realistic noise case, at low dipole moment).

### 4.6 Future Improvements to Increase Laminar Discrimination Accuracy

Future improvements to the methods we describe could further boost laminar discrimination accuracy with low SNR. We used the potential locations of simulation sources as well as some extra vertices as priors for MSP source reconstruction. Future versions of this analysis could use pairs of vertices at corresponding locations on the pial and white matter surfaces as priors. This would take advantage of the sparsity constraints of MSP to determine the most likely location of source activity in each pair (pial or white matter), and the difference between the two could be further amplified by the ROI analysis. Another possibility is to run EBB and MSP source reconstruction in a staged manner, using the results of the EBB source reconstruction as MSP priors. In our simulations, patch size over- or under-estimation significantly affected laminar discrimination accuracy. Intrinsic surface curvature may indicate the extent of lateral connectivity (Ronan et al., 2011), and therefore one possibility for future analyses is to use intrinsic surface curvature to inform patch size estimates. Finally, we found that laminar discrimination was most accurate where the lead field was strongest (Figure 7). Optically pumped magnetometers (OPMs) promise to achieve much higher SNRs than traditional SQUIDs since they can be placed directly on the scalp, boosting the lead field strength (Boto et al., 2017, 2016).

### 4.7 Laminar- and Frequency-Specific Inferences

Theories of brain organization and function such as predictive coding (Bastos et al., 2012; Friston, 2008), communication through coherence (Fries, 2015, 2009, 2005), gating by inhibition (Jensen and Mazaheri, 2010), and communication through nested oscillations (Bonnefond et al., 2017) ascribe distinct roles to deep and superficial cortical laminae as well as low and high frequency oscillations. Support for these theories comes mainly from tract tracing and laminar recordings in nonhuman primates which demonstrate a layer-specific segregation in the origin of feedforward and feedback cortico-cortical connections (Felleman and Van Essen, 1991; Markov et al., 2013) and the distribution of low and high frequency activity (Bollimunta et al., 2011, 2008; Buffalo et al., 2011; Dougherty et al., 2015; Haegens et al., 2015; Maier et al., 2010; Smith et al., 2013; Spaak et al., 2012; van Kerkoerle et al., 2014; Xing et al., 2012). While there is corroborating evidence for these theories in terms of task-modulated frequency-specific activity in the human brain (de Lange et al., 2013; Donner et al., 2009; Jensen et al., 2012; Michalareas et al., 2016; Polanía et al., 2015; Popov et al., 2017), studies linking human oscillatory activity to laminar organization are scarce (Scheeringa et al., 2016). The recent development of high precision MEG with subject-specific head-casts allows non-invasive recording of oscillatory activity in the human brain at previously infeasible SNRs (Meyer et al., 2017; Troebinger et al., 2014b), rendering these theories finally testable in humans (Troebinger et al., 2014a). We have demonstrated that given this high quality data, it is in principle possible to make spatially-localized laminar specific inferences.

## 5 Acknowledgements

JB is funded by a BBSRC research grant (BB/M009645/1). HR is funded by a Wellcome Trust strategic award (104943/Z/14/Z). SM is supported by a Medical Research Council and Engineering and Physical Sciences Research Council grant MR/K6010/86010/1, the Medical Research Council UKMEG Partnership grant MR/K005464/1, and a Wellcome Trust Principal Research Fellowship to Neil Burgess. NA is supported by a Wellcome Trust grant (098352/Z/12/Z). SL was funded by clinical research training grants from the Wellcome Trust (105804/Z/14/Z). The WCHN is supported by a strategic award from Wellcome (091593/Z/10/Z).

## References

Adams, R.A., Shipp, S., Friston, K.J., 2013. Predictions not commands: active inference in the motor system. Brain Struct. Funct. 218, 611–43. doi:10.1007/s00429-012-0475-5

Amir, Y., Harel, M., Malach, R., 1993. Cortical hierarchy reflected in the organization of intrinsic connections in macaque monkey visual cortex. J. Comp. Neurol. 334, 19–46. doi:10.1002/cne.903340103

Arnal, L.H., Giraud, A.-L., 2012. Cortical oscillations and sensory predictions. Trends Cogn. Sci. 16, 390–8. doi:10.1016/j.tics.2012.05.003

Baillet, S., 2017. Magnetoencephalography for brain electrophysiology and imaging. Nat. Neurosci. 20, 327–339. doi:10.1038/nn.4504

Bastos, A.M., Usrey, W.M., Adams, R.A., Mangun, G.R., Fries, P., Friston, K.J., 2012. Canonical microcircuits for predictive coding. Neuron 76, 695–711. doi:10.1016/j.neuron.2012.10.038

Belardinelli, P., Ortiz, E., Barnes, G., Noppeney, U., Preissl, H., 2012. Source reconstruction accuracy of MEG and EEG Bayesian inversion approaches. PLoS One 7, e51985. doi:10.1371/journal.pone.0051985

Bollimunta, A., Chen, Y., Schroeder, C.E., Ding, M., 2008. Neuronal Mechanisms of Cortical Alpha Oscillations in Awake-Behaving Macaques. J. Neurosci. 28.

Bollimunta, A., Mo, J., Schroeder, C.E., Ding, M., 2011. Neuronal mechanisms and attentional modulation of corticothalamic α oscillations. J. Neurosci. Off. J. Soc. Neurosci. 31, 4935–4943. doi:10.1523/JNEUROSCI.5580-10.2011

Bonnefond, M., Kastner, S., Jensen, O., 2017. Communication between Brain Areas Based on Nested Oscillations. eneuro 4, ENEURO.0153–16.2017. doi:10.1523/ENEURO.0153-16.2017

Boto, E., Bowtell, R., Krüger, P., Fromhold, T.M., Morris, P.G., Meyer, S.S., Barnes, G.R., Brookes, M.J., 2016. On the Potential of a New Generation of Magnetometers for MEG: A Beamformer Simulation Study. PLoS One 11, e0157655. doi:10.1371/journal.pone.0157655

Boto, E., Meyer, S.S., Shah, V., Alem, O., Knappe, S., Kruger, P., Fromhold, T.M., Lim, M., Glover, P.M., Morris, P.G., Bowtell, R., Barnes, G.R., Brookes, M.J., 2017. A new generation of magnetoencephalography: Room temperature measurements using optically-pumped magnetometers. Neuroimage 149, 404–414. doi:10.1016/j.neuroimage.2017.01.034

Buffalo, E.A., Fries, P., Landman, R., Buschman, T.J., Desimone, R., 2011. Laminar differences in gamma and alpha coherence in the ventral stream. Proc. Natl. Acad. Sci. U. S. A. 108, 11262–7. doi:10.1073/pnas.1011284108

Bush, P.C., Sejnowski, T.J., 1993. Reduced compartmental models of neocortical pyramidal cells. J. Neurosci. Methods 46, 159–166.

Callaghan, M.F., Josephs, O., Herbst, M., Zaitsev, M., Todd, N., Weiskopf, N., 2015. An evaluation of prospective motion correction (PMC) for high resolution quantitative MRI. Front. Neurosci. 9, 97. doi:10.3389/fnins.2015.00097

Carey, D., Caprini, F., Allen, M., Lutti, A., Weiskopf, N., Rees, G., Callaghan, M.F., Dick, F., 2017. Quantitative MRI Provides Markers Of Intra-, Inter-Regional, And Age-Related Differences In Young Adult Cortical Microstructure. bioRxiv.

Chen, G., Wang, F., Gore, J.C., Roe, A.W., 2013. Layer-specific BOLD activation in awake monkey V1 revealed by ultra-high spatial resolution functional magnetic resonance imaging. Neuroimage 64, 147–155. doi:10.1016/j.neuroimage.2012.08.060

Davatzikos, C., Bryan, N., 1996. Using a deformable surface model to obtain a shape representation of the cortex. IEEE Trans. Med. Imaging 15, 785–795. doi:10.1109/42.544496

De Lange, F.P., Rahnev, D.A., Donner, T.H., Lau, H., 2013. Prestimulus oscillatory activity over motor cortex reflects perceptual expectations. J. Neurosci. 33, 1400–10. doi:10.1523/JNEUROSCI.1094-12.2013

Donner, T.H., Siegel, M., Fries, P., Engel, A.K., 2009. Buildup of choice-predictive activity in human motor cortex during perceptual decision making. Curr. Biol. 19, 1581–5. doi:10.1016/j.cub.2009.07.066

Dougherty, K., Cox, M.A., Ninomiya, T., Leopold, D.A., Maier, A., 2015. Ongoing Alpha Activity in V1 Regulates Visually Driven Spiking Responses. Cereb. Cortex 109, bhv304. doi:10.1093/cercor/bhv304

Elston, G.N., Rosa, M.G., 1997. The occipitoparietal pathway of the macaque monkey: comparison of pyramidal cell morphology in layer III of functionally related cortical visual areas. Cereb. Cortex 7, 432–52. doi:10.1093/CERCOR/7.5.432

Elston, G.N., Tweedale, R., Rosa, M.G., 1999. Cortical integration in the visual system of the macaque monkey: large-scale morphological differences in the pyramidal neurons in the occipital, parietal and temporal lobes. Proceedings. Biol. Sci. 266, 1367–74. doi:10.1098/rspb.1999.0789

Engemann, D.A., Gramfort, A., 2015. Automated model selection in covariance estimation and spatial whitening of MEG and EEG signals. Neuroimage 108, 328–342. doi:10.1016/j.neuroimage.2014.12.040

Felleman, D.J., Van Essen, D.C., 1991. Distributed hierarchical processing in the primate cerebral cortex. Cereb Cortex 1, 1–47.

Fischl, B., 2012. FreeSurfer. Neuroimage 62, 774–781. doi:10.1016/j.neuroimage.2012.01.021

Fischl, B., Dale, A.M., 2000. Measuring the thickness of the human cerebral cortex from magnetic resonance images. Proc. Natl. Acad. Sci. U. S. A. 97, 11050–5. doi:10.1073/pnas.200033797

Fries, P., 2015. Rhythms for Cognition: Communication through Coherence. Neuron 88, 220–235. doi:10.1016/j.neuron.2015.09.034

Fries, P., 2009. Neuronal Gamma-Band Synchronization as a Fundamental Process in Cortical Computation. Annu. Rev. Neurosci. 32, 209–224. doi:10.1146/annurev.neuro.051508.135603

Fries, P., 2005. A mechanism for cognitive dynamics: neuronal communication through neuronal coherence. Trends Cogn. Sci. 9, 474–480. doi:10.1016/j.tics.2005.08.011

Friston, K., 2008. Hierarchical models in the brain. PLoS Comput. Biol. 4, e1000211. doi:10.1371/journal.pcbi.1000211

Friston, K., Harrison, L., Daunizeau, J., Kiebel, S., Phillips, C., Trujillo-Barreto, N., Henson, R., Flandin, G., Mattout, J., 2008. Multiple sparse priors for the M/EEG inverse problem. Neuroimage 39, 1104–1120. doi:10.1016/j.neuroimage.2007.09.048

Friston, K., Mattout, J., Trujillo-Barreto, N., Ashburner, J., Penny, W., 2007. Variational free energy and the Laplace approximation. Neuroimage 34, 220–34. doi:10.1016/j.neuroimage.2006.08.035

Goense, J., Merkle, H., Logothetis, N.K., 2012. High-Resolution fMRI Reveals Laminar Differences in Neurovascular Coupling between Positive and Negative BOLD Responses. Neuron 76, 629–639. doi:10.1016/j.neuron.2012.09.019

Goldenholz, D.M., Ahlfors, S.P., Hämäläinen, M.S., Sharon, D., Ishitobi, M., Vaina, L.M., Stufflebeam, S.M., 2009. Mapping the signal-to-noise-ratios of cortical sources in magnetoencephalography and electroencephalography. Hum. Brain Mapp. 30, 1077–1086. doi:10.1002/hbm.20571

Gramfort, A., Luessi, M., Larson, E., Engemann, D.A., Strohmeier, D., Brodbeck, C., Parkkonen, L., Hämäläinen, M.S., 2014. MNE software for processing MEG and EEG data. Neuroimage 86, 446–460. doi:10.1016/j.neuroimage.2013.10.027

Griffin, L.D., 1994. The Intrinsic Geometry of the Cerebral Cortex. J. Theor. Biol. 166, 261–273. doi:10.1006/jtbi.1994.1024

Guidi, M., Huber, L., Lampe, L., Gauthier, C.J., Möller, H.E., 2016. Lamina-dependent calibrated BOLD response in human primary motor cortex. Neuroimage 141, 250–261. doi:10.1016/j.neuroimage.2016.06.030

Haegens, S., Barczak, A., Musacchia, G., Lipton, M.L., Mehta, A.D., Lakatos, P., Schroeder, C.E., 2015. Laminar Profile and Physiology of the α Rhythm in Primary Visual, Auditory, and Somatosensory Regions of Neocortex. J. Neurosci. 35.

Hämäläinen, M., Ilmoniemi, R., 1984. Interpreting measured magnetic fields of the brain: estimates of current distributions. Tech. rep, Helsinki Univ. Technol.

Hämäläinen, M.S., Ilmoniemi, R.J., 1994. Interpreting magnetic fields of the brain: minimum norm estimates. Med. Biol. Eng. Comput. 32, 35–42. doi:10.1007/BF02512476

Hari, R., Salmelin, R., 1997. Human cortical oscillations: A neuromagnetic view through the skull. Trends Neurosci. doi:10.1016/S0166-2236(96)10065-5

Hillebrand, A., Barnes, G.R., 2011. Practical constraints on estimation of source extent with MEG beamformers. Neuroimage 54, 2732–2740. doi:10.1016/j.neuroimage.2010.10.036

Hillebrand, A., Barnes, G.R., 2003. The use of anatomical constraints with MEG beamformers. Neuroimage 20, 2302–13.

Hillebrand, A., Barnes, G.R., 2002. A quantitative assessment of the sensitivity of whole-head MEG to activity in the adult human cortex. Neuroimage 16, 638–650.

Huber, L., Goense, J., Kennerley, A.J., Trampel, R., Guidi, M., Reimer, E., Ivanov, D., Neef, N., Gauthier, C.J., Turner, R., Möller, H.E., 2015. Cortical lamina-dependent blood volume changes in human brain at 7T. Neuroimage 107, 23–33. doi:10.1016/j.neuroimage.2014.11.046

Hutton, C., Vita, E. De, Ashburner, J., Deichmann, R., Turner, R., 2008. Voxel-based cortical thickness measurements in MRI. Neuroimage 40, 1701. doi:10.1016/j.neuroimage.2008.01.027

Im, K., Lee, J.-M., Yoon, U., Shin, Y.-W., Hong, S.B., Kim, I.Y., Kwon, J.S., Kim, S.I., 2006. Fractal dimension in human cortical surface: Multiple regression analysis with cortical thickness, sulcal depth, and folding area. Hum. Brain Mapp. 27, 994–1003. doi:10.1002/hbm.20238

Jensen, O., Bonnefond, M., Marshall, T.R., Tiesinga, P., 2015. Oscillatory mechanisms of feedforward and feedback visual processing. Trends Neurosci. doi:10.1016/j.tins.2015.02.006

Jensen, O., Bonnefond, M., VanRullen, R., 2012. An oscillatory mechanism for prioritizing salient unattended stimuli. Trends Cogn. Sci. 16, 200–6. doi:10.1016/j.tics.2012.03.002

Jensen, O., Mazaheri, A., 2010. Shaping functional architecture by oscillatory alpha activity: gating by inhibition. Front. Hum. Neurosci. 4, 186. doi:10.3389/fnhum.2010.00186

Jensen, O., Vanni, S., 2002. A New Method to Identify Multiple Sources of Oscillatory Activity from Magnetoencephalographic Data. Neuroimage 15, 568–574. doi:10.1006/nimg.2001.1020

Jones, S.E., Buchbinder, B.R., Aharon, I., 2000. Three-dimensional mapping of cortical thickness using Laplace’s Equation. Hum. Brain Mapp. 11, 12–32. doi:10.1002/1097-0193(200009)11:1<12::AID-HBM20>3.0.CO;2-K

Jones, S.R., Pritchett, D.L., Stufflebeam, S.M., Hämäläinen, M., Moore, C.I., 2007. Neural correlates of tactile detection: a combined magnetoencephalography and biophysically based computational modeling study. J. Neurosci. Off. J. Soc. Neurosci. 27, 10751–10764. doi:10.1523/JNEUROSCI.0482-07.2007

Joshi, S.C., Wang, J., Miller, M.I., Van Essen, D.C., Grenander, U., 1995. Differential geometry of the cortical surface, in: Melter, R.A., Wu, A.Y., Bookstein, F.L., Green, W.D.K. (Eds.), Proc. SPIE 2573, Vision Geometry IV. International Society for Optics and Photonics, pp. 304–311. doi:10.1117/12.216422

Kabani, N., Le Goualher, G., MacDonald, D., Evans, A.C., 2001. Measurement of Cortical Thickness Using an Automated 3-D Algorithm: A Validation Study. Neuroimage 13, 375–380. doi:10.1006/nimg.2000.0652

Kok, P., Bains, L.J., van Mourik, T., Norris, D.G., de Lange, F.P., 2016. Selective Activation of the Deep Layers of the Human Primary Visual Cortex by Top-Down Feedback. Curr. Biol. 26, 371–376. doi:10.1016/j.cub.2015.12.038

Koopmans, P.J., Barth, M., Norris, D.G., 2010. Layer-specific BOLD activation in human V1. Hum. Brain Mapp. 31, 1297–1304. doi:10.1002/hbm.20936

Koopmans, P.J., Barth, M., Orzada, S., Norris, D.G., 2011. Multi-echo fMRI of the cortical laminae in humans at 7T. Neuroimage 56, 1276–1285. doi:10.1016/j.neuroimage.2011.02.042

Kritzer, M.F., Goldman-Rakic, P.S., 1995. Intrinsic circuit organization of the major layers and sublayers of the dorsolateral prefrontal cortex in the rhesus monkey. J. Comp. Neurol. 359, 131–143. doi:10.1002/cne.903590109

Lerch, J.P., Evans, A.C., 2005a. Cortical thickness analysis examined through power analysis and a population simulation. Neuroimage 24, 163–173. doi:10.1016/j.neuroimage.2004.07.045

Lerch, J.P., Evans, A.C., 2005b. Cortical thickness analysis examined through power analysis and a population simulation. Neuroimage 24, 163–173. doi:10.1016/j.neuroimage.2004.07.045

Levitt, J.B., Lewis, D.A., Yoshioka, T., Lund, J.S., 1993. Topography of pyramidal neuron intrinsic connections in macaque monkey prefrontal cortex (areas 9 and 46). J. Comp. Neurol. 338, 360–376. doi:10.1002/cne.903380304

Liuzzi, L., Gascoyne, L.E., Tewarie, P.K., Barratt, E.L., Boto, E., Brookes, M.J., 2016. Optimising experimental design for MEG resting state functional connectivity measurement. Neuroimage. doi:10.1016/j.neuroimage.2016.11.064

López, J.D., Litvak, V., Espinosa, J.J., Friston, K., Barnes, G.R., 2014. Algorithmic procedures for Bayesian MEG/EEG source reconstruction in SPM. Neuroimage 84, 476–487. doi:10.1016/j.neuroimage.2013.09.002

Luders, E., Thompson, P.M., Narr, K.L., Toga, A.W., Jancke, L., Gaser, C., 2006. A curvature-based approach to estimate local gyrification on the cortical surface. Neuroimage 29, 1224–1230. doi:10.1016/j.neuroimage.2005.08.049

Lund, J.S., Yoshioka, T., Levitt, J.B., 1993. Comparison of intrinsic connectivity in different areas of macaque monkey cerebral cortex. Cereb. Cortex 3, 148–62.

Lutti, A., Dick, F., Sereno, M.I., Weiskopf, N., 2014. Using high-resolution quantitative mapping of R1 as an index of cortical myelination. Neuroimage. doi:10.1016/j.neuroimage.2013.06.005

Lutti, A., Hutton, C., Finsterbusch, J., Helms, G., Weiskopf, N., 2010. Optimization and validation of methods for mapping of the radiofrequency transmit field at 3T. Magn. Reson. Med. 64, 229–238. doi:10.1002/mrm.22421

Lutti, A., Stadler, J., Josephs, O., Windischberger, C., Speck, O., Bernarding, J., Hutton, C., Weiskopf, N., 2012. Robust and fast whole brain mapping of the RF transmit field B1 at 7T. PLoS One 7, e32379. doi:10.1371/journal.pone.0032379

MacDonald, D., Kabani, N., Avis, D., Evans, A.C., 2000. Automated 3-D Extraction of Inner and Outer Surfaces of Cerebral Cortex from MRI. Neuroimage 12, 340–356. doi:10.1006/nimg.1999.0534

Maier, A., Adams, G.K., Aura, C., Leopold, D.A., 2010. Distinct Superficial and deep laminar domains of activity in the visual cortex during rest and stimulation. Front. Syst. Neurosci. 4. doi:10.3389/fnsys.2010.00031

Markov, N.T., Ercsey-Ravasz, M., Van Essen, D.C., Knoblauch, K., Toroczkai, Z., Kennedy, H., 2013. Cortical high-density counterstream architectures. Science 342, 1238406. doi:10.1126/science.1238406

McNemar, Q., 1947. Note on the sampling error of the difference between correlated proportions or percentages. Psychometrika 12, 153–157. doi:10.1007/BF02295996

Medvedovsky, M., Taulu, S., Bikmullina, R., Paetau, R., 2007. Artifact and head movement compensation in MEG. Neurol. Neurophysiol. Neurosci. 4.

Meng, X., Rosenthal, R., Rubin, D.B., 1992. Comparing correlated correlation coefficients. Psychol. Bull. 111, 172–175. doi:10.1037/0033-2909.111.1.172

Meyer, H.S., Wimmer, V.C., Oberlaender, M., de Kock, C.P.J., Sakmann, B., Helmstaedter, M., 2010. Number and laminar distribution of neurons in a thalamocortical projection column of rat vibrissal cortex. Cereb. Cortex (New York, N.Y. 1991) 20, 2277–2286. doi:10.1093/cercor/bhq067

Meyer, S.S., Bonaiuto, J., Lim, M., Rossiter, H., Waters, S., Bradbury, D., Bestmann, S., Brookes, M., Callaghan, M.F., Weiskopf, N., Barnes, G.R., 2017. Flexible head-casts for high spatial precision MEG. J. Neurosci. Methods 276, 38–45. doi:10.1016/j.jneumeth.2016.11.009

Michalareas, G., Vezoli, J., van Pelt, S., Schoffelen, J.-M., Kennedy, H., Fries, P., 2016. Alpha-Beta and Gamma Rhythms Subserve Feedback and Feedforward Influences among Human Visual Cortical Areas. Neuron 89, 384–397. doi:10.1016/j.neuron.2015.12.018

Murakami, S., Okada, Y., 2015. Invariance in current dipole moment density across brain structures and species: physiological constraint for neuroimaging. Neuroimage 111, 49–58. doi:10.1016/j.neuroimage.2015.02.003

Murakami, S., Okada, Y., 2006. Contributions of principal neocortical neurons to magnetoencephalography and electroencephalography signals. J. Physiol. 575, 925–936. doi:10.1113/jphysiol.2006.105379

Olman, C.A., Harel, N., Feinberg, D.A., He, S., Zhang, P., Ugurbil, K., Yacoub, E., 2012. Layer-Specific fMRI Reflects Different Neuronal Computations at Different Depths in Human V1. PLoS One 7, e32536. doi:10.1371/journal.pone.0032536

Pascual-Marqui, R., 1999. Review of Methods for Solving the EEG Inverse Problem. Int. J. Bioelectromagn. 1, 75–86.

Pascual-Marqui, R.D., 2002. Standardized low-resolution brain electromagnetic tomography (sLORETA): technical details. Methods Find. Exp. Clin. Pharmacol. 24 Suppl D, 5–12.

Pascual-Marqui, R.D., Michel, C.M., Lehmann, D., 1994. Low resolution electromagnetic tomography: a new method for localizing electrical activity in the brain. Int. J. Psychophysiol. 18, 49–65. doi:10.1016/0167-8760(84)90014-X

Penny, W.D., Stephan, K.E., Daunizeau, J., Rosa, M.J., Friston, K.J., Schofield, T.M., Leff, A.P., 2010. Comparing families of dynamic causal models. PLoS Comput. Biol. 6, e1000709. doi:10.1371/journal.pcbi.1000709

Polanía, R., Moisa, M., Opitz, A., Grueschow, M., Ruff, C.C., 2015. The precision of value-based choices depends causally on fronto-parietal phase coupling. Nat. Commun. 6, 8090. doi:10.1038/ncomms9090

Popov, T., Kastner, S., Jensen, O., 2017. FEF-controlled Alpha Delay Activity Precedes Stimulus-induced Gamma Band Activity in Visual Cortex. J. Neurosci.

Ronan, L., Pienaar, R., Williams, G., Bullmore, E., Crow, T.J., Roberts, N., Jones, P.B., Suckling, J., Fletcher, P.C., 2011. Intrinsic curvature: a marker of millimeter-scale tangential cortico-cortical connectivity? Int. J. Neural Syst. 21, 351–66. doi:10.1142/S0129065711002948

Sakata, S., Harris, K.D., 2009. Laminar structure of spontaneous and sensory-evoked population activity in auditory cortex. Neuron 64, 404–418. doi:10.1016/j.neuron.2009.09.020

Scheeringa, R., Koopmans, P.J., van Mourik, T., Jensen, O., Norris, D.G., 2016. The relationship between oscillatory EEG activity and the laminar-specific BOLD signal. Proc. Natl. Acad. Sci. U. S. A. 113, 6761–6. doi:10.1073/pnas.1522577113

Schubert, D., Kötter, R., Staiger, J.F., 2007. Mapping functional connectivity in barrel-related columns reveals layer- and cell type-specific microcircuits. Brain Struct. Funct. 212, 107–119. doi:10.1007/s00429-007-0147-z

Smith, M.A., Jia, X., Zandvakili, A., Kohn, A., 2013. Laminar dependence of neuronal correlations in visual cortex. J. Neurophysiol. 109, 940–947. doi:10.1152/jn.00846.2012

Spaak, E., Bonnefond, M., Maier, A., Leopold, D.A., Jensen, O., 2012. Layer-specific entrainment of ү-band neural activity by the α rhythm in monkey visual cortex. Curr. Biol. CB 22, 2313–2318. doi:10.1016/j.cub.2012.10.020

Tosun, D., Siddarth, P., Levitt, J., Caplan, R., 2015. Cortical thickness and sulcal depth: insights on development and psychopathology in paediatric epilepsy. Br. J. Psychiatry Open 1.

Troebinger, L., López, J.D., Lutti, A., Bestmann, S., Barnes, G., 2014a. Discrimination of cortical laminae using MEG. Neuroimage 102, 885–893. doi:10.1016/j.neuroimage.2014.07.015

Troebinger, L., López, J.D., Lutti, A., Bradbury, D., Bestmann, S., Barnes, G., 2014b. High precision anatomy for MEG. Neuroimage 86, 583–91. doi:10.1016/j.neuroimage.2013.07.065

Uutela, K., Taulu, S., Hämäläinen, M., 2001. Detecting and Correcting for Head Movements in Neuromagnetic Measurements. Neuroimage 14, 1424–1431. doi:10.1006/nimg.2001.0915

Van Essen, D.C., 2005. A Population-Average, Landmark- and Surface-based (PALS) atlas of human cerebral cortex. Neuroimage 28, 635–662. doi:10.1016/j.neuroimage.2005.06.058

Van Essen, D.C., Drury, H.A., 1997. Structural and Functional Analyses of Human Cerebral Cortex Using a Surface-Based Atlas. J. Neurosci. 17.

Van Kerkoerle, T., Self, M.W., Dagnino, B., Gariel-Mathis, M.-A., Poort, J., van der Togt, C., Roelfsema, P.R., 2014. Alpha and gamma oscillations characterize feedback and feedforward processing in monkey visual cortex. Proc. Natl. Acad. Sci. U. S. A. 111, 14332–14341. doi:10.1073/pnas.1402773111

Wang, X.-J., 2010. Neurophysiological and computational principles of cortical rhythms in cognition. Physiol. Rev. 90, 1195–268. doi:10.1152/physrev.00035.2008

Weiskopf, N., Suckling, J., Williams, G., Correia, M.M., Inkster, B., Tait, R., Ooi, C., Bullmore, E.T., Lutti, A., 2013. Quantitative multi-parameter mapping of R1, PD(*), MT, and R2(*) at 3T: a multi-center validation. Front. Neurosci. 7, 95. doi:10.3389/fnins.2013.00095

Xing, D., Yeh, C.-I., Burns, S., Shapley, R.M., 2012. Laminar analysis of visually evoked activity in the primary visual cortex. Proc. Natl. Acad. Sci. U. S. A. 109, 13871–13876. doi:10.1073/pnas.1201478109

